# Power and shift invariant detection of dynamically coupled networks (PSIICOS) from non-invasive MEG data

**DOI:** 10.1101/129155

**Authors:** A. Ossadtchi, D. Altukhov, K. Jerbi

## Abstract

MEG reflects electrical activity of neuronal assemblies that coalesce and decoalesce in time providing for massively parallel and dynamic flow of information exchange in the brain. There is a growing evidence that our behavior is mediated by simultaneous activity of several interacting and simultaneously active cortical networks. Not only the location of network nodes but also the temporal profiles of such interaction are of interest. The existing methods are primarily concerned with proposing novel measures of synchrony to be applied to the cortical region activity signals extracted from the MEG data using beamforming or other spatial filtering approaches.

In this work we propose to render the task of estimating network topology and temporal profiles as a source estimation problem but in the space of interacting topographies. The proposed point of view has a number of benefits as it allows to use the wealth of techniques and intuition accumulated in the community when dealing with source estimation problems. Operating in the interacting topographies space we first propose subspace projection method to significantly reduce the spatial leakage effect in the cross-spectral matrix. We then use subspace matching metrics to extract a set of networks with their synchronicity profiles that explain the variance in the sensor space cross-spectral matrix, much like regular dipoles and their activations explain the variance in the regular evoked responses.

The preliminary results of application of this method to a simple self-paced voluntary finger movement paradigm comparing allowed to observe functional coupling between primary motor and supplementary motor area and establish that this coupling can be characterized with near-zero phase delay between the oscillatory activity of these two regions. This observed synchrony pattern is physiologically plausible and is in agreement with the current hypothesis about the neural mechanics of the motor acts.

## 1 Introduction

The rapidly growing field of brain connectomics provides compelling evidence for a prominent role of functional and structural connectivity in mediating healthy brain function (REFS). It is now widely accepted that behavior is mediated to a great extent by the coordinated activity of interacting cortical areas (e.g. Varela et al. [2001], Baker et al. [2014], Ossadtchi et al. [2010], Bastin et al. [2017], Rodriguez et al. [2012], Jerbi et al. [2007]). Established research points to a prominent role of brain networks (van den Heuvel et al. [2012]) and neuronal rhythms (Buzsaki G. 2006, Schnitzler and Gross [2005]) in providing mechanistic processes and dynamic brain-wide architectures. Collectively, such networks of oscillatory brain patterns provide a rich repertoire of neuronal communication paths necessary for cognition (Fries [2015]).

Beyond the spatial structure of a network, which is given by its nodes and edges, the time-varying and frequency-dependent strength of the interactions are critical; They are thought to capture the dynamic nature of such aggregations that coalesce and de-coalesce over time reflecting the complex structure of information flow in the brain. Therefore, the identification and characterization of the spatial, temporal and spectral patterns of such interactions through electrophysiology is necessary in order to foster novel insights into the basic mechanisms and functional role of brain connectivity (Varela et al. [2001]). Despite important contributions from neuroimaging modalities, such as functional magnetic resonance imaging (fMRI), the detailed fine-grained dynamics are only amenable to electrophysiological measurement techniques such as electroencephalography (EEG), intracranial EEG and magnetoencephalography (MEG). When combined with source estimation techniques, MEG offers a unique combination of sub-centimeter spatial accuracy with millisecond-range temporal resolution (Hamalainen et al. [1993], Baillet et al. 2001, Gross et al. [2013]).

Considerable effort has been directed towards developing metrics and methodological frameworks to increase the diversity and reliability of cortical interaction measures. The large number of methods implemented and used over the last decades cover a wide range of approaches ranging from standard time and frequency-domain interaction metrics (such as correlation and coherence) to measures that quantify specific dimensions of signal coupling, such as phase-based or band-limited amplitude relationships, directionality, cross-frequency interactions or non-linear associations (Bastos and Schoffelen [2016], Marzetti et al. [2008], Schoffelen and Gross 2009, Colclough et al. [2016], Colclough et al. 2015, Greenblatt et al. [2012], Kaminski and Blinowska [2014], Hillebrand and Stam [2014], Nolte et al. [2004b], Lachaux et al. [1999], O’Neill et al., Brookes et al. 2012, Brookes et al. [2011], Hillebrand et al. [2012], Hipp et al. [2012], Vinck et al. [2011], Stam et al. 2007, Chella et al. [2016], Chella et al. [2015], Soto et al. [2016], Wibral et al. [2011], Ioannides et al. [2000]).

One of the most prominent challenges that faces non-invasive MEG/EEG connectivity estimation is arguably the spatial leakage effect which significantly complicates non-invasive studies of connectivity. In 2004, Nolte et al. (Nolte et al. [2004a]) suggested the use of the imaginary part of coherency as solution to the spatial leakage issue; by ignoring the real component, imaginary coherence is de facto insensitive to zero-phase interactions. Since signal leakage is instantaneous, this method guarantees that any detected coupling reflects true physiological interaction. This was followed by further innovative measures based solely on the imaginary part of the cross-spectrum such as the Phase Lag Index (PLI) (Stam et al. 2007) and its weighted version (wPLI) (Vinck et al. [2011]). These and other related metrics such as orthogonalized envelope correlation (Hipp et al. [2012], Colclough et al. 2015) have now been widely adopted. Yet, despite providing an important pragmatic step forward, these metrics can underestimate true coupling because they are -by construction-blind to true physiological zero-phase interactions, if any are present among the signals.

Another often overlooked limitation arises from the fact that, by definition, the distribution of magnitudes of the imaginary part of the cross-spectrum of coupled sources is centered on the sine of the mean phase difference. As a result, in cases where the coupled sources have near-zero time lags, coupling measures exploiting the imaginary part of cross-spectrum are tuned to a low SNR signal. In other words, not only are these metrics blind to zero-phase lag interactions, but their sensitivity is weak for phase lags approaching zero. Ironically, in many cases, empirically observed neuronal synchrony is often characterized by a very small phase lag between the timeseries of coupled neuronal ensembles (e.g. Roelfsema et al. [1997]). Indeed, true physiological zero-lag and near-zero lag coupling patterns may have several explanations. They could be the result of the presence of the third source providing common influence on the time series of the two generators, or alternatively, the result of bidirectional interaction between neuronal assemblies. Macro-scale analysis shows that two cortical areas engaged in bidirectional interaction are likely to generate near-zero phase-lag synchrony as a result of reciprocal interaction or positively correlated common input (Rajagovindan and Ding [2008]). Moreover, near-zero lag coupling scenario may also be linked to the effect of detuning of synchronized populations with slightly different dynamical properties in order to adapt to a "single and global rhythm" (Pikovsky et al. 2001; Schuster and Wagner [1989]). Therefore, in order to map cortical networks in healthy subjects under a wide range of experimental conditions and exercise freedom in fine-tuning the experimental designs it is important to develop synchrony detection techniques which are unaffected by source leakage (MEG field spread, or EEG volume conduction) but which are nevertheless capable of reliably identifying leakage-free zero-phase interactions and reliably detecting near-zero phase lagged coupling. This is precisely the aim of the present study, where we derive a new framework for the non-invasive estimation of connectivity with lag-invariant sensitivity.

Most MEG and EEG data synchrony analysis methods start with estimating cortical source activation timeseries, which are subsequently used for source-space analysis of coupling. This approach leads to propagation of timeseries estimation errors to the synchrony metrics used. The cross-talk is only one of the problems that researchers face when using such two-step procedures. While this approach might be inevitable when probing non-linear interaction effects, linear interactions can be studied using promising and intuitive approaches based on utilization of the generative model of sensor space cross-spectrum. As a matter of fact, an approach exploiting the spatial structure of the imaginary part of the observed cross-spectrum via subspace correlation metrics has recently been developed by Ewald et al. [2014]). However, as discussed above, the use of the imaginary part of the cross-spectrum may entail weaker sensitivity for near-zero lag coupling, compared to larger phase lags.

In the new method proposed here, we start with the generative model of the observed cross-spectrum of sensor signals and utilizes the fact that the sensor space cross-spectrum is a linear combination of the outer product of interacting source topographies. The coefficients of such linear combinations are source-space cross-spectrum values. We then consider the vectorized form of this model and demonstate that a simple projection operation defined in the product space of sensor signals can be used to alleviate the effects of spatial leakage (SL) on the real part of the cross-spectrum. Next, we examine the properties of the suggested procedure by the means of Monte-Carlo simulations and we show that such debiasing of the real part of the cross-spectrum also delivers better sensitivity to near zero lag coupling compared to the methods exploiting only the imaginary part of the cross-spectrum. We then analyze MEG datasets of 10 subjects involved in the auditory language task and demonstrate that implementation of the suggested projection operation to remove the undesired SL effect with the use of real-part of the cross-spectrum furnishes significantly better reproducibility of sensor-space and source-space synchrony analysis results as compared to the methods that rely solely on the imaginary part of the cross-spectrum. Based on the above and given very modest computational requirements of the method we conclude that the described technique overcomes previous limitations by providing a reliable interaction metric with a sensitivity profile that does not depend on the value of the phase lag. We believe that the proposed method, dubbed PSIICOS (Phase shift invariant imaging of coherent sources), can become a useful addition to the corpus of existing approaches for inter-areal connectivity estimation in MEG and EEG data.

## 2 Methods

### 2.1 MEEG signal model and preliminaries

MEEG data recorded by a *K-* sensor array can be written as the following linear combination

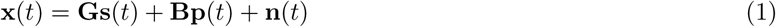

In other words, multichannel signal at each instance of time, **x**(*t*) is a noisy additive mixture of source topographies **G** = [**g**_1_*, …,* **g**_*L*_] weighted by the corresponding activation timeseries **s**(*t*) = [*s*_1_(*t*)*, …, s*_*L*_(*t*)] of the dipolar sources, along with a task-unrelated contribution from the sources with topographies **B** = [**b**_1_*, …,* **b**_*M*_] with task-unrelated activations **p**(*t*) = [*p*_1_(*t*)*, …, p*_*M*_ (*t*)], and an additive random noise vector **n** (*t*). Topographies of the task-related sources form an *R*-dimensional task-relevant signal subspace and topographies of task unrelated sources form an *L*-dimensional coherent interference subspace.

MEG and EEG recorded brain activity cab be described as a non-stationary mixture narrow-band components and is best characterized in time-frequency domain. Let **X**(*t, f*), **S**(*t, f*), **P**(*t, f*) represent time-frequency transform coefficients of **x**(*t*), **s**(*t*) and **p**(*t*) correspondingly obtained via some kind of a linear time-frequency transform.

Let us consider the expression for the sensor-space cross-spectral matrix, the sufficient statistics for estimation of the source-space linear coupling. By computing the expectation of the outer product **X**(*t, f*)**X**^*H*^ (*t, f*) we obtain

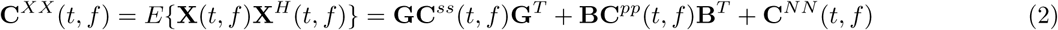

Here **C**^*NN*^ is a diagonal cross-spectral matrix of the spatially incoherent additive sensor noise, **C**^*ss*^(*t, f*) = *E* {**S**(*t, f*)**S**^*H*^ (*t, f*)} is a non-diagonal source-space cross-spectral matrix that we need to estimate and **C**^*pp*^(*t, f*) = *E* {**P**(*t, f*)**P**^*H*^ (*t, f*)} is the source-space cross-spectral matrix of the task unrelated brain noise sources. The generic formulation given by (2) is valid both for task-based and task-free spontaneous recordings; First, in the case of a task-based paradigm, the coherence of the brain noise sources is by definition not supposed to be related to the task and therefore **C**^*pp*^(*t, f*) is a diagonal matrix. Note that although task independent inter-areal coupling may exist, the key aspect here is that they will not be systematically be modulated by the task, or else they would be captured by the term **C**^*ss*^(*t, f*). Second, in the spontaneous data case, the brain-noise represented by the forward matrix **B** is taken as a superposition of incoherent sources and therefore **C**^*pp*^(*t, f*) is also a diagonal matrix. In other words, here also, all resting-state couplings will be captured by the term **C**^*ss*^(*t, f*). Furthermore, in the case of epoched data the approximation of expectation operation is computed by averaging the time-frequency coefficients over trials. When dealing with spontaneous data (e.g. resting state) the mean operation can be computed over some contiguous time intervals or a set of such intervals with similar spatial statistics as illustrated in the recent work Baker et al. [2014].

Next, let us expand matrix multiplications in the right-hand side of equation (2) to obtain

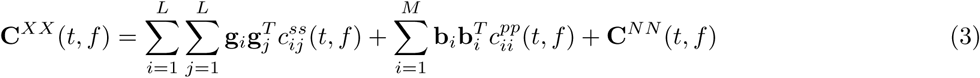

This equation is in fact, a linear generative model for the sensor space cross-spectral matrix. **C**^*XX*^ (*t, f*) is a linear combination of rank one elementary network topography matrices 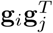 and 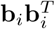 with weights *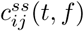* and 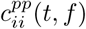. These time-frequency dependent weights are split into two groups. The first group of weights *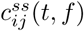* are the cross-spectral coefficients for the *i*-th and the *j*-th source. In case *i* == *j* this coefficient is real and reflects the power of the *i*-th source, if *i* ≠ *j* the weights are in general complex-valued cross-spectral coefficients of the source activation timesereis. The second group of weights 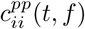 reflects the power of incoherent sources with activations *p*_*i*_(*t*), see (1).

In other words, equation 3 states that the sensor-space crossspectral matrix is an element of *K*^2^ dimensional product space and can be represented as a superposition of elementary 2-node network "sources" with "topographies" 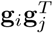 and complex-valued activations 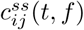. The observed mixture is contaminated by the incoherent brain-noise sources via 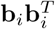 "topographies". Motivated by the structural similarity of equations 1 and 3 we will refer to 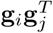 as network topograhies and correspondingly 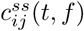 as temporal synchrony profiles.

In general **C**^*XX*^ (*t, f*) is complex. Since 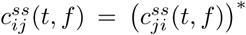 for the real-valued source timeseries we can separate the real and imaginary parts of **C**^*XX*^ (*t, f*) in equation (3) and obtain

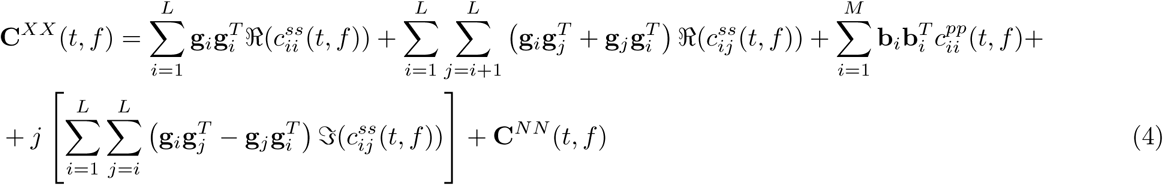

As we can see from (4) the real part of the sensor-space crossspectrum contains contributions not only from the off-diagonal elements of the source-space crossspectrum matrix 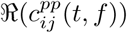 but also from its diagonal elements 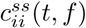 reflecting the power of the coupled sources. Additionally, the power of mutually incoherent brain noise sources *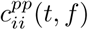* also contributes to the real part of the cross-spectrum.

By contrast, the imaginary part of the cross-spectrum is only sensitive to the non-instantaneously coupled sources because ℑ (*c*_*ij*_(*t, f*)) = 0 for the instantaneous coupling. In order to emphasize the fact that the imaginary part of the cross-spectrum is immune to (i.e. unaffected by) spatial leakage effects, we intentionally kept the notation *j* = *i* (instead of *j* = *i* + 1) in the second sum of the last double summation term in (4). We can see that for the *i -th* source, this term does not make any contribution to the imaginary part because the corresponding elementary network topography matrix 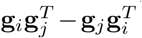 vanishes when *i* = *j*. This fact was used in Nolte et al. [2004a] to introduce the imaginary part of coherence as a linear coupling measure immune to spatial leakage. This seminal study was followed by several related approaches and extensions such as PLI (Stam et al. 2007) and wPLI (Vinck et al. [2011]) all solely exploiting the imaginary part of the crosss-pectrum.

Interpreting 4 from a linear algebra point of view, we can also say that the subspace of the real part of the crossspectral matrices is spanned by 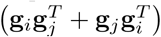 matrices and the corresponding complex valued component subspace is spanned by 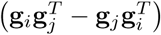 matrices. The latter fact was for instance used in Ewald et al. [2014] to introduce the Wedge-MUSIC approach, which aims to uncover coherent sources based on the spatial structure of the imaginary part of the sensor-space crossspectrum.

The discussed spatial mapping effect is only one of two elements that condition the signal-to-noise ratio (SNR) of the observed sensor space activity generated by a 2-node network source with topography 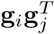 and complex time-frequency activation profiles *c*_*ij*_(*t, f*). The distribution of the magnitude between the real and imaginary parts of the source-space cross-spectral coefficient *c*_*ij*_(*t, f*) depends on the mean phase difference Δ*ϕ* of the timeseries of the coupled sources. The magnitude of the imaginary part is proportional to *sin*(Δ*ϕ*) while the magnitude of the real-part follows a *cos*(Δ*ϕ*) profile. As a result, when the mean phase difference is close to *pi/*2 the magnitude of the imaginary part of the cross-spectrum achieves its highest value yielding an SNR which facilitates the detectability of such coupling based on the imaginary component of the sensor-space cross-spectrum. However, when Δ*ϕ* is small (i.e. approaches zero), the magnitude of the imaginary part of the cross-spectrum drops drastically which results in low SNR and thereby poor detectability of coupling between such sources, when uniquely based on the imaginary part of the cross-spectrum. And, as described above, the use of the real part of the sensor-space cross-spectrum is hindered by the fact that the latter is severely affected by the spatial-leakage effect. In summary, while previous techniques based on the imaginary part of the cross-spectrum do rule out that the detected interactions are entirely explained by spatial leakage, they have an important limitation: their sensitivity profile varies as a function of the phase lag, from high sensitivity at mean Δ*ϕ* = *pi/*2, and gradually decreasing sensitivity which ultimately vanishes at Δ*ϕ* = 0). The goal of the present study is to derive an interaction estimation technique with a sensitivity profile that does not change as a function of the phase shift (i.e. phase-shift invariant)

### 2.2 Vectorized form of cross-spectrum

The ultimate goal of synchrony studies is to estimate the elements *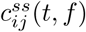*, *i* ≠ *j* of the source space crossspectral matrix for specific combination of time and frequency values. Tensor-like structure of equations (3) and (4) obscures the fact that they can be managed using the standard and efficient vector linear algebra framework and that synchrony estimation problem can be formally cast in exactly the same way as the standard source estimation problem in EMEG but in the *K*^2^ dimensional product space.

To illustrate this, let us first consider the vectorized version of equation (4) by unfolding the source-space cross-spectral matrix and its additive components into *K*^2^ dimensional vectors. This operation is called vectorization and has the standard *vec*() notation. This linear operation of vectorization puts us in the *K*^2^-dimensional linear space of vectors and allows to think about the problem of detection of coupled sources and estimatbion the time-frequency profiles of their coupling strength in much the same way as we think about locating active sources from regular EMEG data and estimating their time-frequency activation patterns. Additionally, as we will show in the following sections this approach leads to a straightforward and intuitive way to remove the spatial leakage effect and remain invariant to the mean phase difference (Δ*ϕ*) of activity of coupled sources.

Without loss of generality, let us consider the vectorized form of the cross-spectral matrix corresponding to some predefined sub-band specified by central frequency *f*_0_. This leaves us with only variability over time and the vectorized cross-spectrum will simply be a matrix of dimension *K*^2^*×T* where *T* is the number of time samples. Therefore the vectorized expression of the time-varying cross-spectrum at frequency *f*_0_ will look as follows

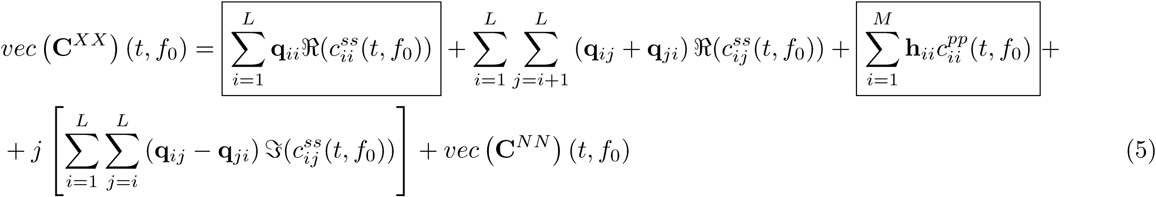

with 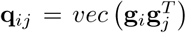 and 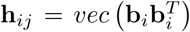. In the above 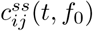 denote the source space cross-spectrum value at *f*_0_ between the activity of the *i*-th and the *j*-th source at time *t*. In what follows for compactness we will drop the *f*_0_ symbol.

Thus, (5) is the generative model equation and therefore the unknown time-varying source-space cross-spectrum elements can be estimated by a variety of well known techniques. The inherent temporal variability comes in naturally and is treated in exactly the same way as dipole source timeseries are treated in the EMEG source estimation task. However, in equation (5) the role of a source is played by a pair of interacting dipoles with topographies expressed in *K*^2^ dimensional product space. The observed sensor space cross-spectrum is simply a noisy time-varying linear combination of interacting topographies with coefficients being unknown source space cross-spectral values.

### 2.3 Spatial leakage effect

Consider equation (5) and note that the framed terms 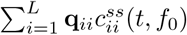 and 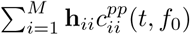 represent spatial leakage (SL) of source power into the observed sensor-space cross-spectrum. The vectorised sensor-space cross-spectrum *vec* **C**^*XX*^ (*t, f*_0_) receives rank-one component **q**_*ii*_ in the amount proportional to the power of the *i*-th source 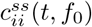 at frequency *f*_0_. The incoherent sources also contribute to the vectorised cross-spectrum as 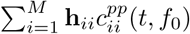.

Therefore, we can say that the spatial leakage subspace *S*_*SL*_ of the vectorised cross-spectrum is spanned by vectors **q**_*ii*_*, i* = 1*,.., M*. Similarly note that the real-part of the source space cross-spectrum modulates power in the *S*_*ℛ*_ = *span*(**q**_*ij*_ +**q**_*ji*_) subspace and the imaginary part of the source-space cross-spectrum fills in *S*_*ℑ*_ = *span*(**q**_*ij*_*-***q**_*ji*_) subspace.

It is straightforward to show that *S*_*ℛ*_*⊥ S*_*ℑ*_, *S*_*ℑ*_*⊥ S*_*SL*_ and that *S* _*ℛ*_*nS*_*SL*_ *≠ø*. The diagram in Figure [1] illustrates this interrelation between the vector components that comprise the sensor-space cross-spectrum. Physically, the intersection of *S*_*ℛ*_ and *S*_*SL*_ is primarily occupied by the contributions from the real-part of cross-spectrum of functionally coupled sources located close to each other whose activations can be characterized by a low value of mean phase difference.

**Figure 1:**
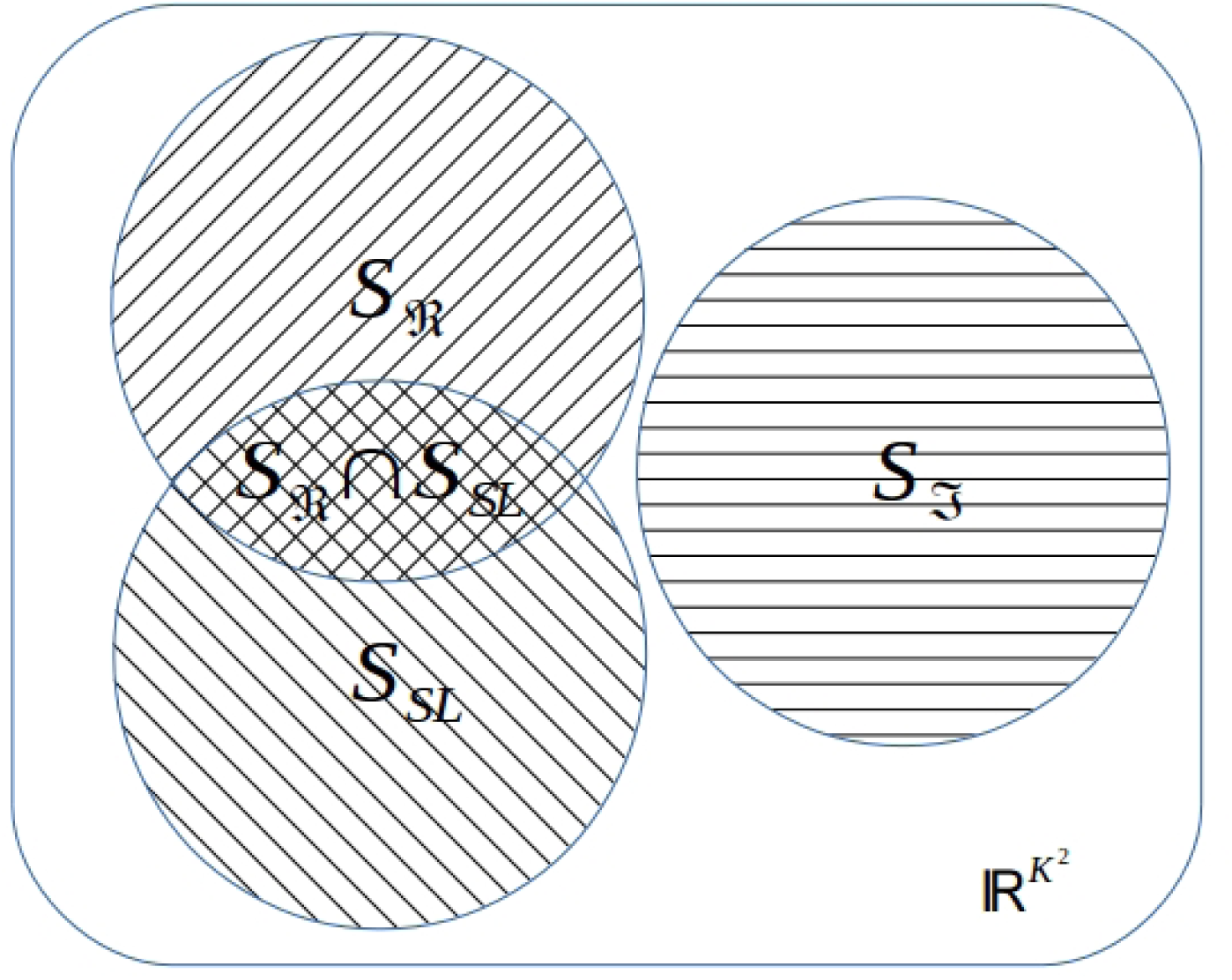
Interrelation of subspaces of the sensor space cross-spectrum. The spatial leakage subspace and the real part subspace overlap and are orthogonal to the subspace of the imaginary component of the cross-spectrum. The intersection of the two former subspaces contains contributions from the spatial leakage and from the real-part of actual cross-spectrum of sources located close to each other whose activity can be characterised by a low value of mean phase difference.

### 2.4 Removing spatial leakage effect

Given the linear mixture equation (5), in order to remove the contribution of the spatial leakage on the real part of the cross-spectrum we need to remove the first and the fourth element of the rhs of equation (5).

We suggest a general procedure that does not require estimation of source power from the particular dataset at hands. We will simply project the vectorised cross-spectrum away from the subspace *S*_*SL*_ spanned by **q**_*ii*_*, i* = 1*, …, K*. As outlined above this procedure will not leave *S* intact and therefore we will need to achieve some balance between the extent to which we mitigate the SL and how much the *S* subspace is affected. We will manage this trade-off by adjusting the rank of our projection operator built as follows.

We will first form the *K*^2^ *× L* matrix **F** of SL topographies **q**_*ii*_*, i* = 1*,.., L* as

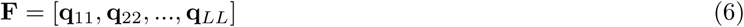

In order to limit the projecting rank *R* and at the same time capture the maximum energy contained in the *S*_*SL*_ subspace we will perform the SVD of **F** as **F** = **USV**^*T*^ and form matrix **U**_*R*_ of the first *R* left singular vectors **U**_*R*_ = [**u**_1_, **u**_2_*, …,* **u**_*R*_] whose columns span the most powerful subspace of *S*_*SL*_. The extent to which the spatial leakage component is removed depends solely on the geometric properties of the head-sensor array system. Therefore, the informed choice of a value for R can be done based on the plots similar to 4 showing the relative attenuation of the *S*_*SL*_ and *S*_*Re*_ components.

We then form matrix 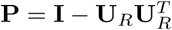 and use it to create almost SL free vectorised cross-spectrum as

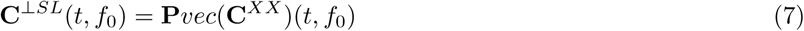

Note, that we created **P** based on the entire set of source space elements (as represented in the forward model matrix **G**) and therefore we addressed the SL contribution of all sources including that of incoherent sources in the model (1).

### 2.5 Accommodation of arbitrary orientation

Anatomically, the dipole orientations coincide with the direction of apical dendrites of the pyramidal neurons and are therefore appear to be predominantly orthogonal to the cortical mantle. Modern tools of MRI data analysis allow for a very accurate extraction and precise parametrization of the cortical surface with number of nodes on the order to tens of thousands, which in turn results into reasonable accuracy of orientation specification. The uncertainty that remains may be efficiently modeled with such techniques (Hamalainen 2006 loose orientation).

Because of memory and processing time limitations when performing exploratory source-space synchrony analysis we have to use a significantly downsampled version of the cortex that contains only several thousands of nodes. Such downsampling drastically reduces the requirements to computational resources but introduces significant uncertainty in orientations of elementary sources.

Therefore, a common practice become to restrict the source space to the downsampled cortical mantle and to model the arbitrary orientations of node dipoles by representing each elementary source with the triplet of orthogonal dipoles. In case of MEG, since the magnetic field outside the spherical conductor produced by a dipole with radial orientation, the triplet can be efficiently replaced by a pair of dipoles in the tangential plane calculated for each node.

For an arbitrary orientation vector at some *i*-th vertex *θ* = [*θ*^*x*^*θ*^*y*^]^*T*^ the corresponding dipole topography is 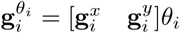 where 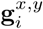 are the topographies of the two orthogonally oriented dipoles in the tangential plane at the *i*-th vertex. Varying the orientation angle we will obtain an infinite set of SL topography vectors 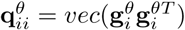. It is easy to show that in case of two dimensional tangent space (as we have in MEG) all these vectors will belong to the three dimensional subspace 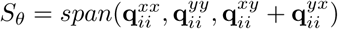.

Therefore, in order to accommodate the arbitrary orientation constraint equation 6 has to be replaced by

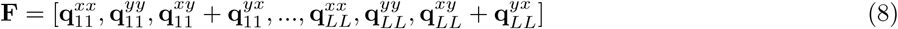

The projection matrix is then found as described in section 2.4.

### 2.6 Source space analysis

We used equations (4) and (5) in order to understand and derive the projector away from the SL subspace. Once we have this projector we can return to the much more compact equation (3) and after applying the projection operation write

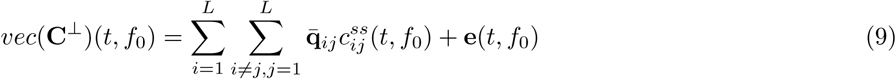

The projected interaction topography vectors 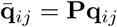 and the noise term **e**(*t, f*) contain the leftovers of the spatial leakage component from the sources of interest and incoherent brain-noise that survived the rank-truncated projection away from the SL subspace.

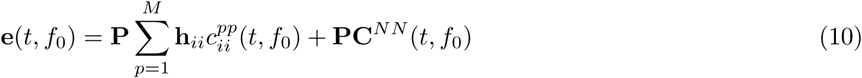

b Consideration of the vectorized form of the cross-spectral matrix allowed us to construct the efficient projection procedure for strongly reducing the SL components from the real-part of the sensor-space cross-spectrum. Additional benefits of the vectorization based view is that it allows us to appreciate the fact that equation (9) is simply a linear regression equation similar to the one we use when solving the source estimation problem but formulated in the product space of sensor signals. Instead of source activation time-series (as in (1)) or their time-frequency profiles in the regular equation the unknown regression coefficients in equation (9) are the time-frequency profiles of the source space cross-spectral coefficients.

The goal here is to estimate 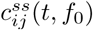 which can be accomplished by a plethora of methods developed for solving the classical inverse problem in MEG and EEG. In this work in order to demosntrate the performance of the suggested projection procedure and perform ROC analysis based on the simulated source-space ground truth we resort onto the simplest possible inverse mapping procedure based on projecting the data onto the interacton topographies vector. We perform a simple scan by computing the inner product of the corresponding projected interaction topographies 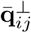 at each time slice of the vectorized and projected cross-spectrum matrix *vec*(**C**^*⊥*^(*t, f*_0_)) to obtain *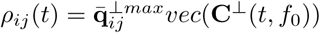*

We calculate the following quantity 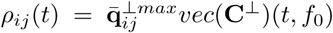 that uses 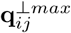 interaction topography that maximizes the norm of *ρ*_*ij*_(*t*) over all possible orientations of the *i*-th and the *j*-th source. Since after the projection *vec*(**C**^*⊥*^)(*t, f*_0_) practically does not contain the contribution from the SL subspace we can use **q**_*ij*_ instead of its projected version 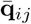. Then, with very little degradation in performance we can optimize over orientations efficiently by computing the largest singular value of a 2 *×* 2 matrix 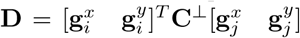. The left **u**_*max*_ and right **v**_*max*_ singular vectors corresponding to this largest singular value determine the orientation of the two interacting dipoles. Based on the magnitude of *ρ*_*ij*_(*t*) we can judge about the presence of interacting sources at coordinates corresponding to the *i*-th and *j*-th node at time instance *t*.

## 3 Results

In the three numerical studies described below we used the 3-shell sphere based forward model matrices calculated using the Brainstorm software exploiting the Freesurfer extracted cortical surfaces with 15000 vertices. At each node of the cortical mesh we modeled a pair of orthogonal dipole sources located in the locally tangential plane.

### 3.1 Effect of projection on the source space cross-spectrum components

As evident from the diagram in Figure [1] because of the orthogonality of *S* and *S*_*SL*_ subspaces the projection operation does not alter the spatial structure of the imaginary part of the cross-spectrum. On the contrary, the components from *S* subspace carrying information about the real part of source-space cross-spectrum do get affected by such a projection operation and by removing the SL variance from the sensor-space cross-spectrum we will inevitably remove some part of the variance that comes from the real component of the source space cross-spectrum.

Clearly, the attenuation of the variance in the *S*_ℛ_ subspace is most pronounced for sources with correlated topographies. To explore the dependence of attenuation on the correlation coefficient between the topographies of the coupled sources for each combination of network node indexes *i* and *j* (*i ≠ j*) we calculated the norms of the product-space topographies 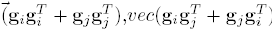 and 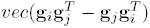 spanning the three subspaces *S*_*SL*_, *S*_*ℑ*_ and *S*_ℛ_ of the sensor-space cross-spectrum, see figure [1]. We did so before the projection and after the projection operation and presented the results in figure [2] using the scatter plot. For each two-node network with nodes in the *i*-th and the *j*-th locations of the cortical surface we plot three points color-coded using the three different (red,blue,brown) palettes. The gradations of the blue color is used to depict the norm of the product-space topography vector **q**_*ij*_ + **q**_*ji*_ spanning the *S*_ℛ_ subspace of the sensor-space cross-spectrum. The gradations of the red color is used to depict the norm of the product-space topography vector **q**_*ij*_*-***q**_*ji*_ spanning the *S*_*ℑ*_ subspace of the sensor-space cross-spectrum. Finally, the gradation of the brown color is used to depict the norm of the product-space topography vector **q**_*ii*_ + **q**_*jj*_ spanning the *S*_*SL*_ subspace of the sensor-space cross-spectrum under the assumption of equal power of the *i*-th and *j*-th source. The position of each colored point along the x-axis is determined by the correlation coefficient of topographies **g**_*i*_ and **g**_*j*_ of the dipolar sources comprising the elementary network. We have also plotted the same data but used source grid node distance for the x-axis, see figure [3]. Color saturation reflects the density of the scattered points.

**Figure 2:**
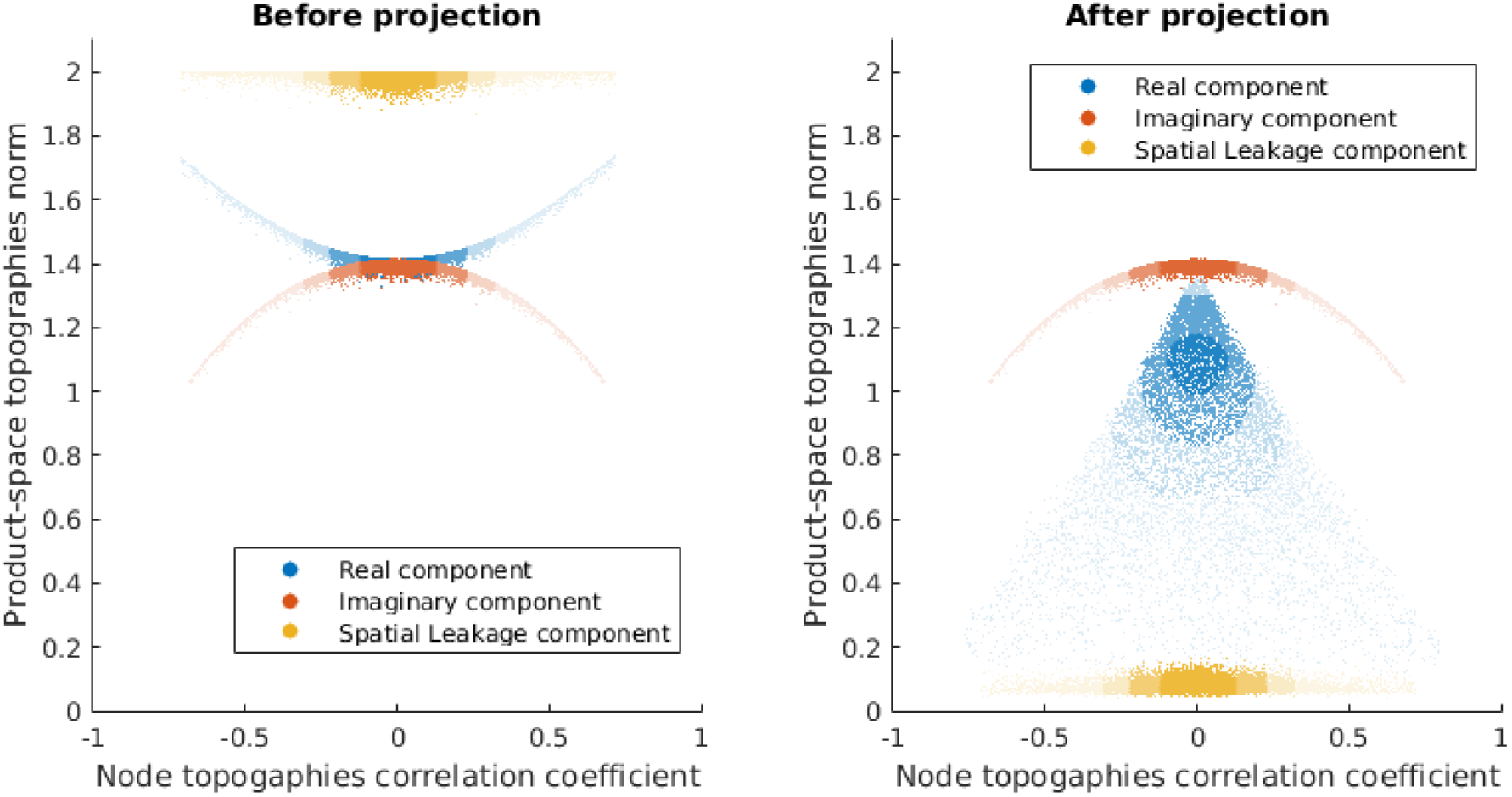
Product-space topography norms for the three subspaces of the sensor-space cross-spectrum before (left panel) and after (right panel) the projection as a function of the correlation coefficient of network node source topographies. Before the projection (left panel) the sensor-space cross-spectrum is dominated by the source power component (brown). After the projection the manifestation of the source power on the sensors gets reduced by more than a factor of 25 on average. We are also witnessing the inevitable but significantly less dramatic attenuation (1.6 is the mean value) of power in the *S*_ℛ_ component of the sensor-space cross-spectrum.

**Figure 3:**
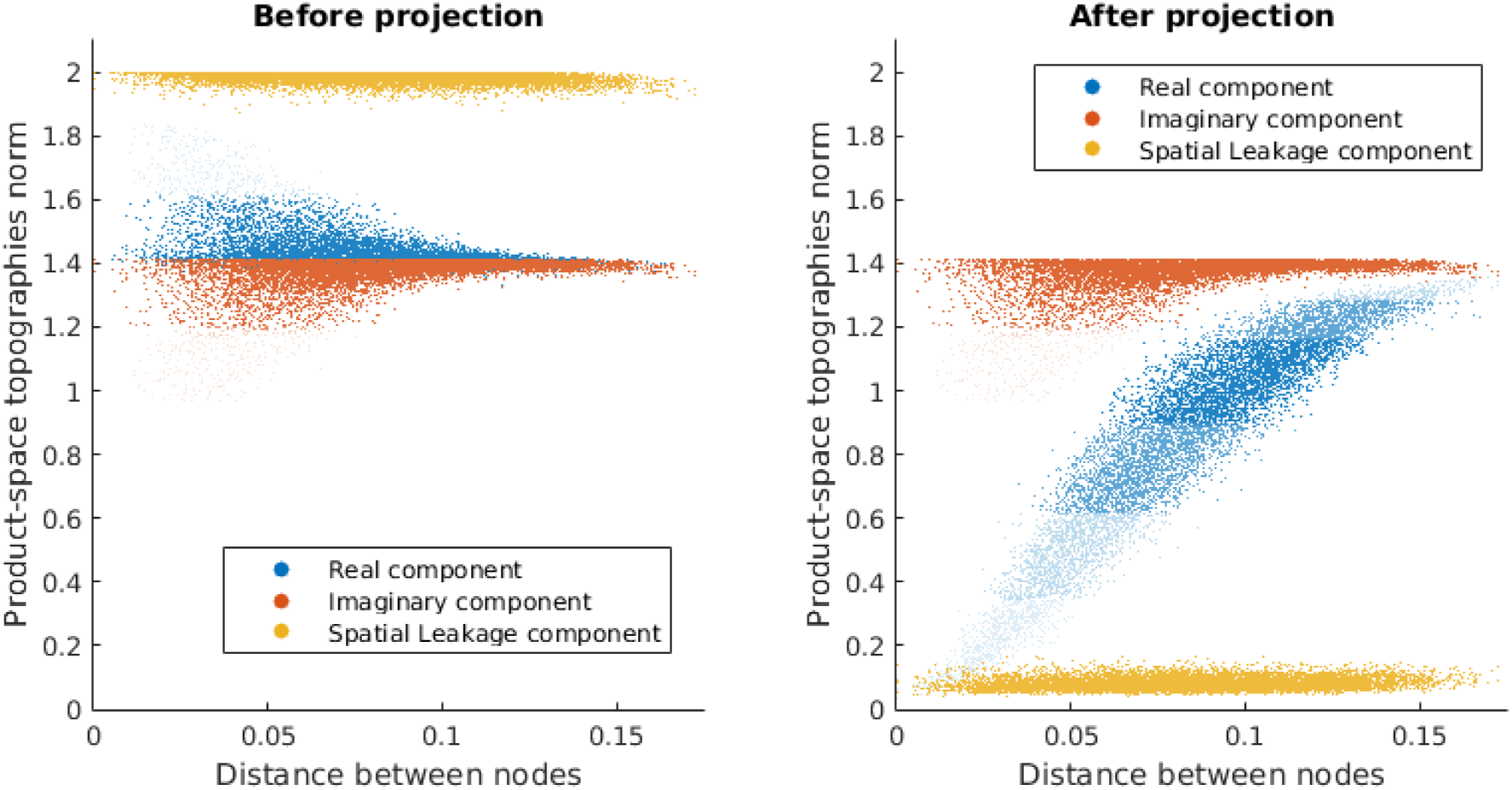
Product-space topography norms for the three subspaces of the sensor-space cross-spectrum before (left panel) and after (right panel) the projection as a function of the distance between the pair of elementary network nodes

As we can see from the diagram on the left panels of 2,3 before the projection operation the SL component (brown) dominates the sensor space cross-spectrum. After applying the described projection operation of rank 500 (right panel) we can observe that the significant reduction in the SL component power for all values of the dipole topographies correlation coefficient. Because the *S*_*SL*_ and *S*_ℛ_ subspaces intersect the projection procedure also reduces the power in the *S*_ℛ_ subspace as it can be seen by comparing the blue scatter in the left and the right panels. However, the reduction in the norm of the product-space topography vectors spanning *S*_ℛ_ is significantly less than the attenuation suffered by the SL subspace topography vectors.

We have also studied the dependence of the attenuation factors in all three subspaces as a function of the projection rank. Figure [4] shows the average attenuation of the power in the three subspaces as a function of the projection rank. To obtain this plot we performed a Monte-Carlo analysis. At each iteration we randomly selected a subset of 200 sources and calculated all the the vectors from the three subspaces *S*_*SL*_*, S*_ℛ_*, S*_ℑ_. We used sources with fixed orientations orthogonal to the cortical mantle. We then varied the rank of the projection and computed the projected versions of these vectors. To quantify the attenuation effect we calculated the average ratio of the original to the projected vector norms for each value of the projection rank. We repeated this process 100 times and averaged the result.

**Figure 4:**
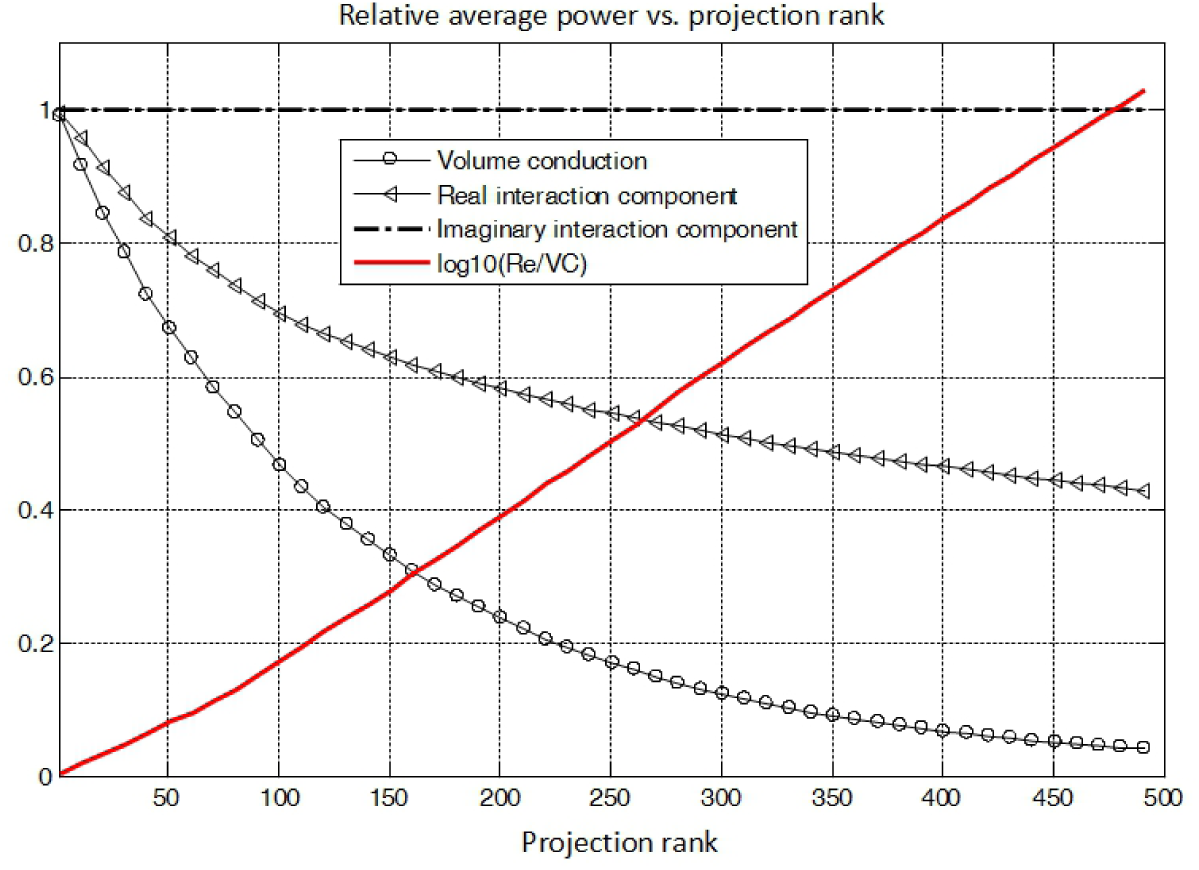
Attenuation of variance in the three subspaces as a function of projection rank

Because of the intersection of the *S*_*SL*_ and *S*_ℛ_ subspaces the projection operation attempting to suppress the SL component power inevitably leads to suppression of power in the *S*_ℛ_ subspace. The less power in the *S*_ℛ_ subspace is suppressed for the fixed SL power suppression factor the better the performance is. Therefore, in addition to the attenuation curves for the subspaces we have also plotted the log-ratio of attenuation coefficients observed for the vectors in the *S*_ℛ_ subspace to that for the *S*_*SL*_ subspace. The use of the SVD operation to form the projection operator allowed for the observed significantly faster reduction in the spatial leakage subspace *S*_*SL*_ power than in the *S*_ℛ_ subspace. In other words we see that the increased projection rank leads to a greater dominance of power in *S* subspace over that in the *S*_*SL*_ subspace.

We have thus shown that cross-spectrum **C**^⊥^(*t, f*_0_) the projected away from the SL subspace is dominated by the energy in the true connectivity subspace. The product-space topographies of the interacting pairs can be easily computed as 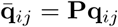 and the corresponding generating model of the projected cross-spectrum (see equation 9) can be used to perform inference on the source-space cross-spectral coefficients 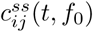.

### 3.2 Forward model inaccuracy effect

The described projection procedure requires forward model (FM) operator that in a realistic setting inevitably comes with noise. In this section we study the performance of the the proposed projection operation in presence of the structured and non-structured inaccuracies in the forward model.

We used two models of noise in this study. The first model corresponded to spatially unstructured noise and was implemented simply by adding the appropriately scaled matrix random noise matrix to the FM model operator matrix. We sampled form the *N* (0, 1) distribution to generate a FM noise realization matrix. We then scaled this by multiplying its elements by the square-root of the mean trace value of matrix **GG**^*T*^ where **G** is the forward model matrix. We then added this matrix to the actual(true) forward model and adjusted the amount of noise by parameter *α*.

The second model corresponds to a more realistic scenario of spatially structured distortion. To generate the spatially structured noise matrix we used the head models computed for *N* = 10 subjects and calculated the pairwise differences between the forward models for each pair of subjects. We then computed the structured noise matrix as the average of these pair-wise differences. We then standardized the resulting structured noise matrix added it to the true forward model. As in the previous case we adjusted the amount of noise by parameter *α* used to scale the noise matrix.

Potentially, the inaccuracies in the FM may cause the simultaneous reduction in the SL power attenuation and the decrease of *S*_ℛ_ subspace power. We performed Monte-Carlo simulations to numerically study these effects. At each iteration we randomly selected a subset of 200 sources and calculated mean attenuation ratio for the three subspaces. We did so for both structured and unstructured noise cases. The results are shown in figure [4]. We used two different projection rank values (350 and 500) in this study. The left panel shows the dependence of the SL attenuation factor as a function of the FM noise intensity *α*. On the right panel we can see the attenuation factor for *S*_ℛ_ subspace variance. While the inaccuracies in the forward model do cause deterioration of performance of the proposed projection scheme, for typical MEG FM noise levels of 10 % () we have almost the same value of the SL power attenuation and only 10 % additional decrease in the *S* subspace variance.

**Figure 5:**
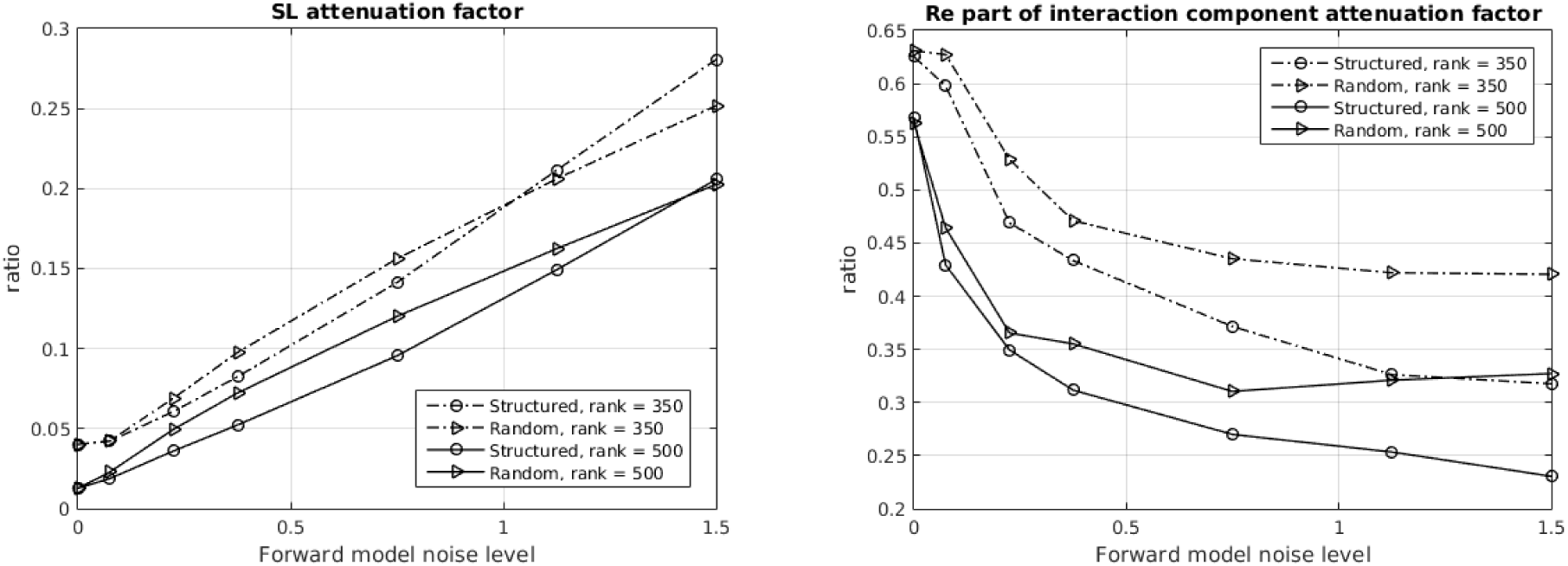
Effects of FM noise on the projection performance. The left panel shows the dependence of SL attenuation factor as a function of the FM noise intensity *α*. On the right panel we can see the attenuation factor for the variance in the *S*_ℛ_ subspace. While the inaccuracies in the forward model do cause deterioration of performance of the proposed projection scheme, for typical MEG FM noise levels of 10 % () we have almost the same value of the SL power attenuation and only 10 % additional decrease in the *S*_ℛ_ subspace variance.

In this section we have described our numerical studies investigating the deterioration of performance of the posed projection scheme due to the forward model inaccuracies. Based on the presented numerical results we conclude that the method is sufficiently robust to tolerate typical FM modeling inaccuracies and can be used in a realistic setting. However, as with many other model based techniques (e.g. beamforming) the efforts of creating more accurate forward models will have a tangible pay-off and therefore need not be neglected.

### 3.3 Simulation setup

In order to compare the proposed technique against other existing methods we ran a set of realistic simulation studies. We used Freesurfer extracted cortical surface with 15000 vertices and calculated the forward model matrix **G**^*HR*^ with two columns per vertex corresponding to the two orthogonally oriented dipoles in the local tangential plane. We simulated ERP experiment data with 100 repetitions (epochs) of the task. The induced activity of coupled sources was modeled with two 10 Hz sinusoidal functions with random phase w.r.t. to stimulus onset but probabilistically connected via phase difference term *δϕ* sampled from a random distribution taking values in the [*-π/*4*, π/*4] range. To simulate the transient nature of such a network within each trial we modulated this synchronized activity by a window function *w*(*t*). Temporal profiles of activity of the three networks and their spatial topologies are shown in figure [6]. Network nodes were chosen to coincide with one of the vertices of the high density grid with 15000 nodes. The corresponding pair of columns of the forward matrix **G**^*HR*^ was summed with scaling to reflect the orthogonal to the cortex orientation and then used to project source activation into the sensor space. Note, that in order to simulate the real-life circumstances when source coordinates do not necessarily match a vertex in a mesh we used the high resolution cortex only to simulate the data. We employed the 10 times sparser representation of the cortex with 1503 node for analysis of the simulated datasets.

**Figure 6:**
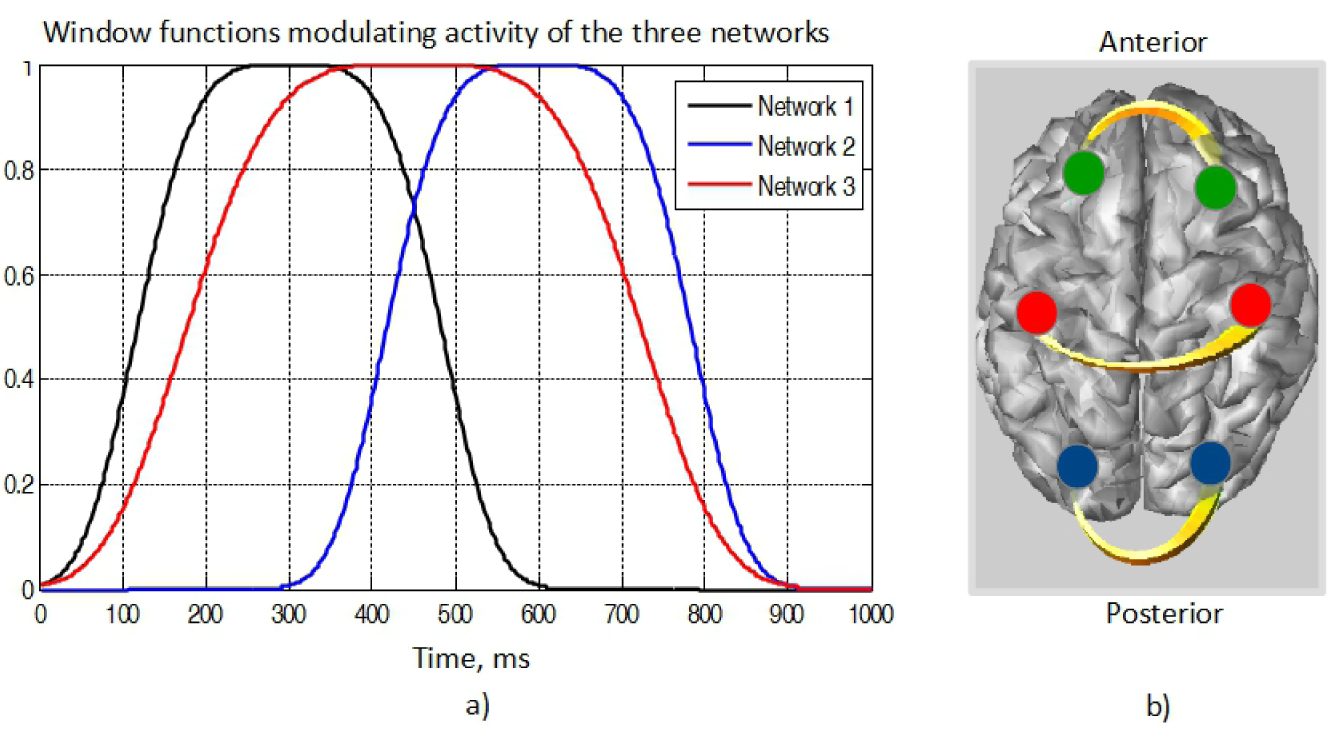
Three simulated networks. a) -temporal profiles of activity of each of the three networks. b) -spatial topography of the simulated networks.

We modeled brain noise with *Q* = 1000 spatially coherent, task-unrelated cerebral sources whose locations and time series varied with each realization. Source locations matched nodes of the high resolution cortical surface (15000 vertices). The activation time series were narrow-band signals obtained via zero-phase filtering of realizations of Gaussian (pseudo)random process by the fifth order band-pass IIR filters in the bands corresponding to theta (4-7 Hz), alpha (8-12 Hz), beta (15-30 Hz) and gamma (30-50 Hz, 50-70 Hz) activity. Their relative contributions were scaled in accordance with 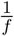 characteristic of the realistic EMEG spectrum. We scaled the brain noise components to match typical signal-to-noise ratio of real-life recordings. To project these sources into the sensor space the corresponding columns of **G**^*HR*^ were linearly combined to represent the orthogonally oriented dipole in each of the *Q* chosen nodes. We simulated 100 epoch ERP data and for each epoch a new randomly picked set of noisy sources was chosen and new noisy timeseries were generated.

During the separate Monte-Carlo studies we report first, we used only one pair of interacting sources placed at random locations on the high resolution cortex at each MC trial. The modulating window function coincided with the depicted in figure [6] and activated the network only during the first half of each trial. At each MC trial we generated a 100 epoch dataset with fixed spatial configuration of target sources and varying from epoch to epoch constellation of brain noise sources. For detection of networks we used the entire trial time range to mimic the situation when we don′t know the interval during which the network is active. For each MC trial we then computed frequency specific sensor space cross-spectral matrix by averaging over epochs the outer product of channel timeseries Fourier coefficients that fall in the [8 − 12*Hz*] frequency range. For each MC trial we calculated a ROC curve then averaged these ROC curves over the entire set of 100 Monte-Carlo trials. Since the proposed method is designed to achieve uniform performance for all values of the phase-lag between the coupled sources we have performed separate studies for two different phase-lag values: *ϕ* = *π/*2 −*π/*20 and *ϕ* = *π/*20 radians.

### 3.4 Performance metrics used

As a threshold-free performance metrics we used the Precision-Recall and ROC curves. The Precision-Recall metrics suites well the situation when we have a small number of true positives and the number of choices is large. The ROC curve, *sensitivity* vs 1 *-specif icity* plot, in this case is informative only for extremely high specificity values. Precision (or positive predictive value), recall (sensitivity) and specificity are defined as follows

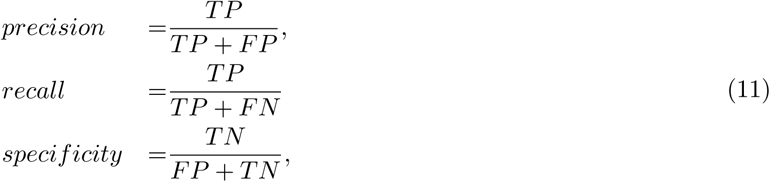

where TP, FN,FP and TN stand for true positives, false negative, false positive and true negative detections correspondingly. In order to calculate these quantities we need to label all possible connections between the source-grid nodes as those that belong and those that not to the simulated networks. Since for simulating the data we used a high resolution cortical mesh with 15000 grid nodes and for detection step we used a 10 times sparser version of the cortical mesh we employed the notion of *δ*-cylinder. Each *n*-th true network is defined in our simulations by a pair of nodes with coordinates 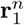 and 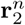. For each such node we define a set of indices of cortical mesh nodes 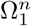 and 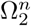 whose coordinates fall into the *δ* neighborhood of 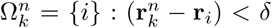 and 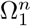, i.e. 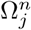 for *k* = 1, 2.

Then, a connection between a pair of nodes from 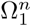 and 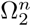 subsets (but not within one subset) is considered as a true connection corresponding to network *n*.

As we can see, from the last line of equation (11), since all node-pairs that fall within 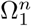 and 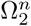 subsets constitute a small fraction of all possible connections and TN is extremely large only high *specif icity* values are of interest. Since the ROC curve shows *sensitity* vs 1 *specif icty* we will be primarily interested in behavior of this curve for low abscissa values. In our plots we considered ROC curves in [0, 0.01] domain. At the same time, the precision metrics operates only within the subsets 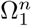 and 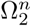 which makes the precision-recall plots a convenient performance metric in our scenario.

Precision-recall(P-R) curve clearly reflects the performance in a single network case. For multiple true networks (which is the case in the second set of simulations) the high area under the P-R curve may be obtained even if only one out of several true networks is detected. In order for the P-R plot to adequately reflect the performance we encoded with dye size the number of true networks the detected connections come from.

### 3.5 Comparative performance study

In this section we will describe our results comparing the proposed PSIICOS technique against three other methods for detection of synchronous sources. One of the most popular methods is the Dynamic Imaging of Coherent Sources (DICS) technique developed by J. Gross. The method boils down to applying a frequency domain beamformer to the Fourier coefficients of the multichannel sensor-space data followed by computing the coherence values of thus extracted Fourier domain representation of the source space signals. The orientations of the two sources are chosen to maximize the source-space cross-spectrum magnitude value. In the original formulation this method appears to be quite sensitive to the Spatial Leakage effects. We thus modified this technique and used only the imaginary part of the source-space cross-spectral matrix. We refer to this modified DICS as iDICS technique. The third method we compare PSSICOS against is Geometric Correction Scheme (GCS) approach developed by xxxx. This method is conceptually most closely related to the proposed technique here and suggests to apply a model based geometric transformation to the second source data to remove the spatial leakage effect from the first source. As in the original paper we use the MNE as the inverse operator and to compute the MNE weights we have supplied the MNE with generally unknown sensor space brain noise covariance matrix. This information is not required by PSIICOS.

#### 3.5.1 Monte-Carlo study of receiver operating characteristics

To study the improvement in detection characteristics due to afforded by PSIICOS taking into account the SL free real part of the cross-spectrum and appreciate the extent of such improvements for various spatial configuration of networks we performed a Monte-Carlo study according to the principles described in the section 3.3. We considered two different values of band specific single trial noise level and two different phase lag values, *π/*20 and *π/*2 −*π/*20. For each combination of noise level (1 and 0.2) and phase lag parameters (*π/*20 and *π/*2*−π/*20) we performed 1000 MC trials. For each MC trial we randomly picked a pair of source space grid nodes to represent the coupled sources (elementary network). We then generated 100 trials of ERP data with fuzzy phase coupling of the 10 Hz osciallations. Each trial was 1 second long and the coupling took place only during the first 500 ms of each trial. To mimic the real-life situation when we don′t know upfront the time range the coupling takes place at we used the entire trial duration to compute the sensor space cross-spectral matrix. At each MC trial we compute the Precision-recall and ROC curves and averaged them over 1000 MC trials.

As we can see from the plot in figure [7] for each of the simulation conditions PSIICOS consistently outperforms the other methods and allows to achieve reasonable performance invariant to the phase lag value. As judged by the ROC curve for phase lag of *ϕ* = *π/*2 *−π/*20 the iDICS delivers performance comparable to that of PSIICOS. For *ϕ* = *π/*20 the iDICS fails to adequately detect the networks due to significantly decreased SNR of representation of nearly zero-phase lag coupling in the imaginary part of the cross-spectrum. At the same time the PSIICOS technique manages to retain the performance for both SNR values. The reason for such an improved performance lies in the use of the proposed projection operation that frees the real part of the cross-spectrum from the undesired SL component leaving only the component modulated by the real part of the cross-spectral coefficient of the two sources. For near zero phase lag this very components captures the information about the coupling and therefore PSIICOS is able to detect it.

**Figure 7:**
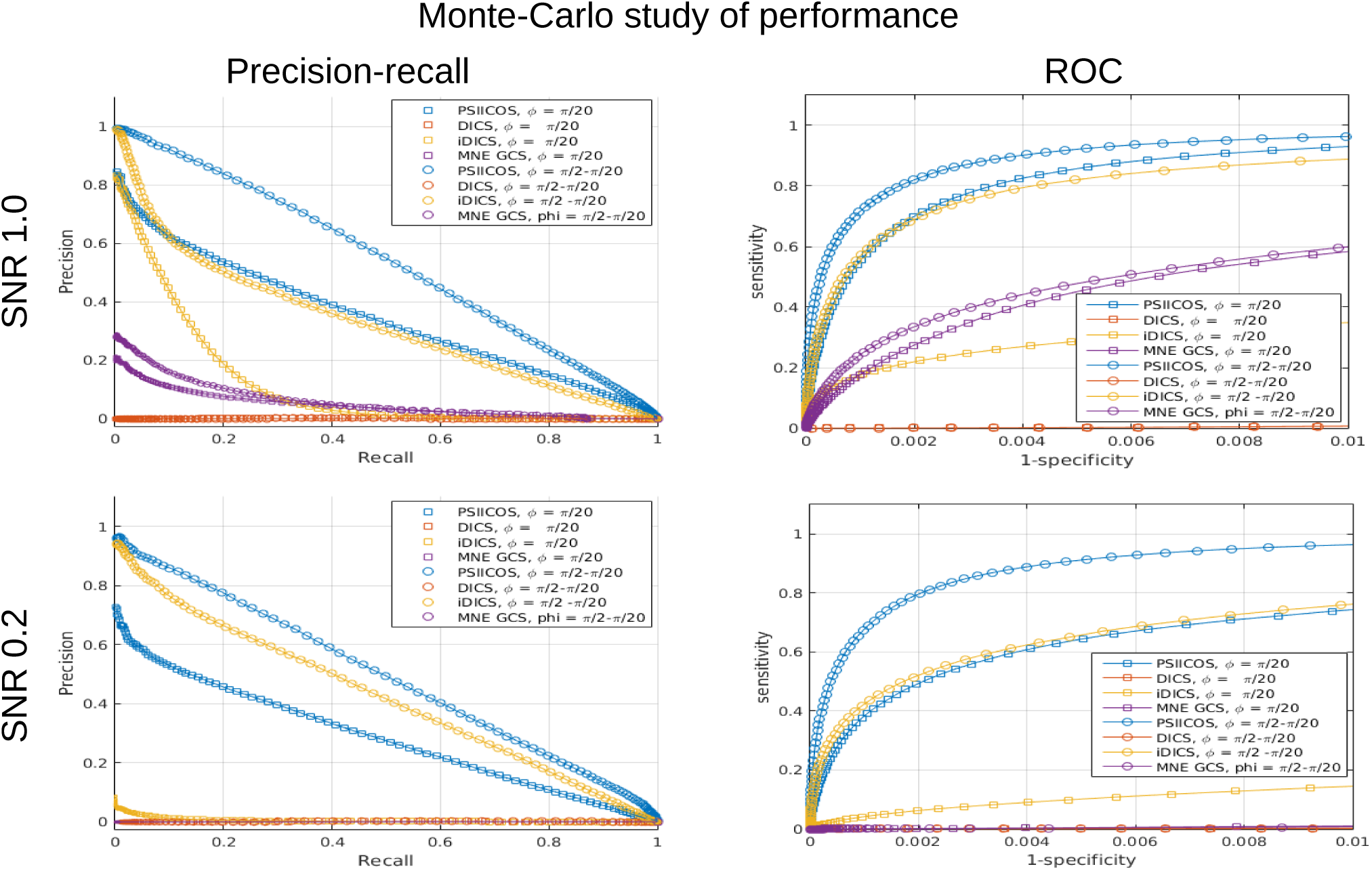
Precision-recall and ROC curves comparing the network detection performance of the proposed approach vs DIC, iDICS and GCS MNE techniques for two different SNR values based on 1000 Monte-Carlo trials.

However, for the lower SNR case we observe a greater gap between PSICCOS′s performance for two different phase lag values. This can be due to the presence in MC trials networks with nodes located close to each other so that the real part of the real-interaction term gets significantly affected by the projection operation, see figure [4]. The iDICS technique, however, practically stops operating in these conditions for near zero lag coupling and low SNR scenario.

#### 3.6 Realistic simulations

##### 3.6.1 Projection procedure illustration

In this section we simulated three networks whose activity overlapped in time as it is illustrated in figure [6]. We have filtered the simulated data in the 8-12 Hz band and calculated the time varying cross-spectral tensor by averaging over trials the outer product of Hilbert transform coefficients of the data at at each time slice. We thus obtained matrix **C^XX^**, see equation (3) which is ready to be used in PSIICOS analysis pipeline. We perform these simulations in the heavy noise conditions and the real part of the cross-spectrum is originally very seriously contaminated as it can be seen from the two left panels of figure [8] for SNR = 0.2 scenario. We would also like to note here that unlike in the single network case the SNR here is computed for all three networks simultaneously and therefore given a significant overlap of activity of these networks in time the actual SNR of each network is even lower.

**Figure 8:**
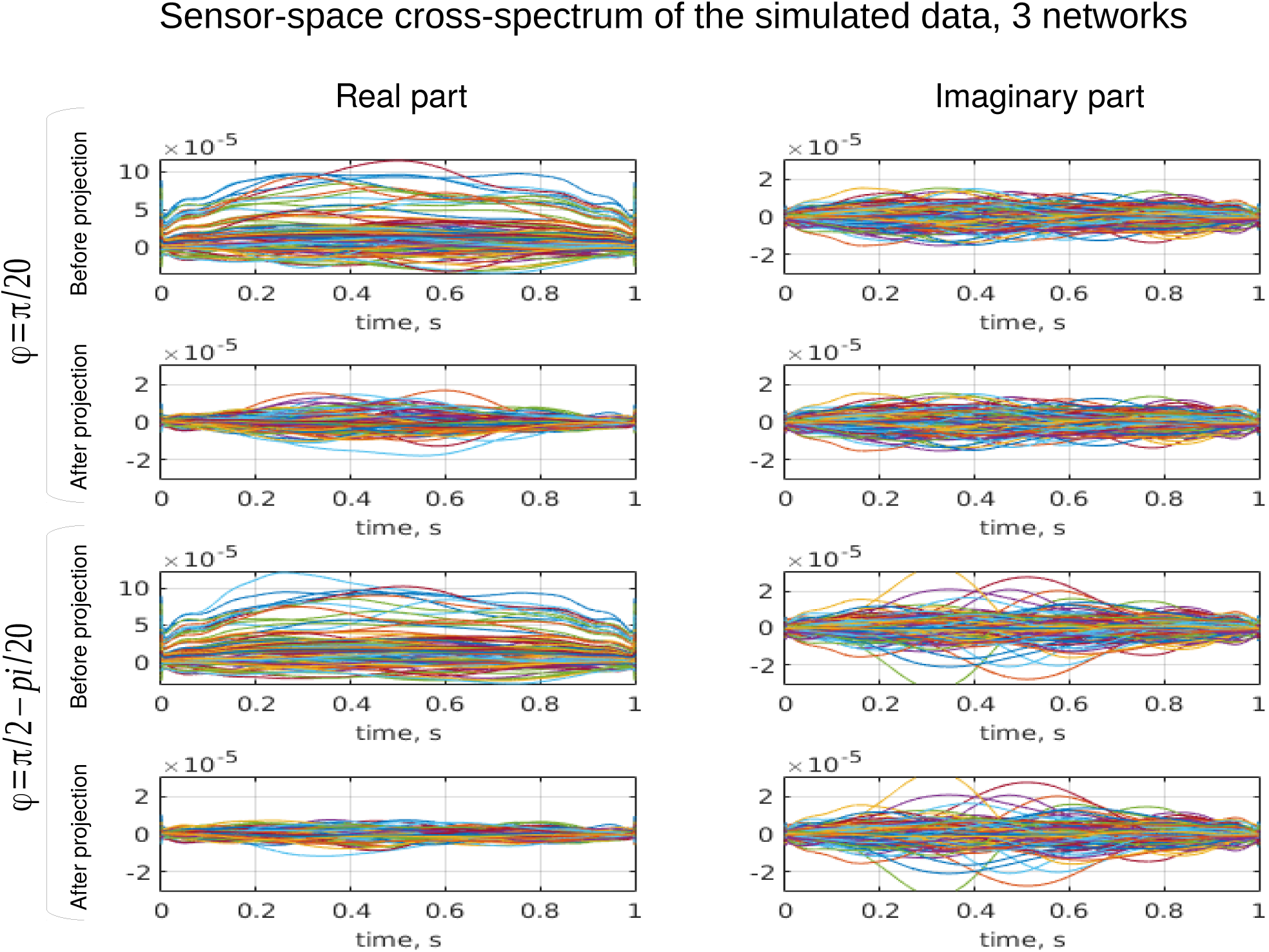
To illustrate the effects of the proposed projection procedure we are showing simulated sensor-space time-resolved cross-spectra for SNR = 0.2 case and two phase lag values before and after the projection. The top half of the plots corresponds to near zero phase lag *ϕ* = *π/*20 and the bottom half for the phase lag close to *π/*2

The top half of figure [8] shows the time resolved cross-spectral matrices and the result of applying the projection procedure for nearly zero phase-lag case. In this case almost all of the information about the coupling is contained in the real part of the cross-spectrum and is significantly contaminated by the SL component. This can be appreciated by comparing this top left plot to the third from the top left plot showing the real part of the cross-spectrum but for the nearly *π/*2 phase lag so that all the coupling related variance is shifted to the imaginary component leaving primarily the SL contribution in the real part. The second from the top left plot shows that the application of the projection operation removes the undesired SL contribution so that the simulated temporal dynamics of the three networks (see figure [6]) can be seen even in the sensor space data. Also, note the scale change between the plots showing the original and the projected cross-spectra to appreciate the amount of noise present in the data. as expected the imaginary part of the cross-spectrum is not affected by the projection operation. For the nearly *π/*2 phase lag the imaginary part of the cross-spectrum contains recognizable (see figure [6]) temporal dynamics leaving the real part almost empty after the projection (bottom left plot).

Next in figure [9] we are showing the source-space spatial structure of the simulated data scan depicting top 0.1% of connections with the highest value of the source-space connectivity statistics for PSIICOS and three other methods. As one can see only the PSIICOS technique retains reasonable operation for all range of the studied conditions. Consistent with our previous results the iDICS technique performance nearly pars that of PSIICOS in the close to *π/*2 phase lag case. The MNE GCS technique for the high SNR case quite reliably picks the two frontal networks for both values of the phase lag but completely stops operating for the low SNR and near zeros phase lag case. For the close to *π/*2 phase lag case under the low SNR it picks up only the middle network (the one with the highest individual SNR) and generates a large number of false positives.

**Figure 9:**
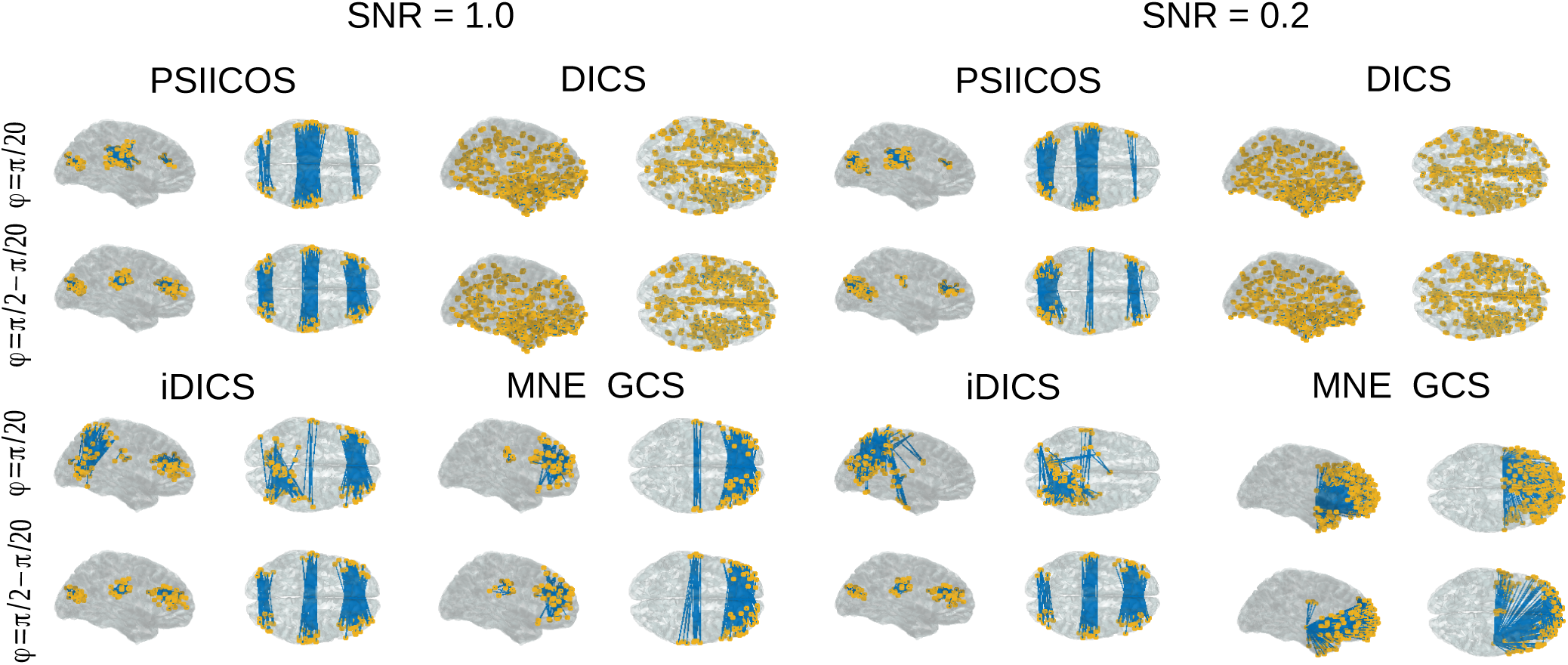
Spatial structure of source-space scan of the simulated data depicting top 0.1% of connections with the highest value of the source-space connectivity statistics for PSIICOS and three other methods. As one can see only the PSIICOS techniques retains reasonable operation for all range of the studied conditions. Consistent with our previous results the iDICS technique performance nearly pars that of PSIICOS in the close to *π/*2 phase lag case. The MNE GCS technique for the high SNR case quite reliably picks the two frontal networks for both values of the phase lag but completely stops operating for the low SNR and near zeros phase lag case. For the close to *π/*2 phase lag case under the low SNR it picks up only the middle network (the one with the highest individual SNR) and generates a large number of false positives.

Further, to systematically illustrate the comparative analysis of the four techniques considered in this paper we are showing in figure [10] the modified precision-recall plots where as described earlier we encoded with dye size the number of true networks the detected connections come from. The left half of the plots shows the entire precision-recall plot and the right panel focuses on the 0-0.15 range. As we can see qualitatively the picture corresponds to the conclusions made based on the visual analysis of the spatial structure of detected connections depicted in figure [9].

**Figure 10:**
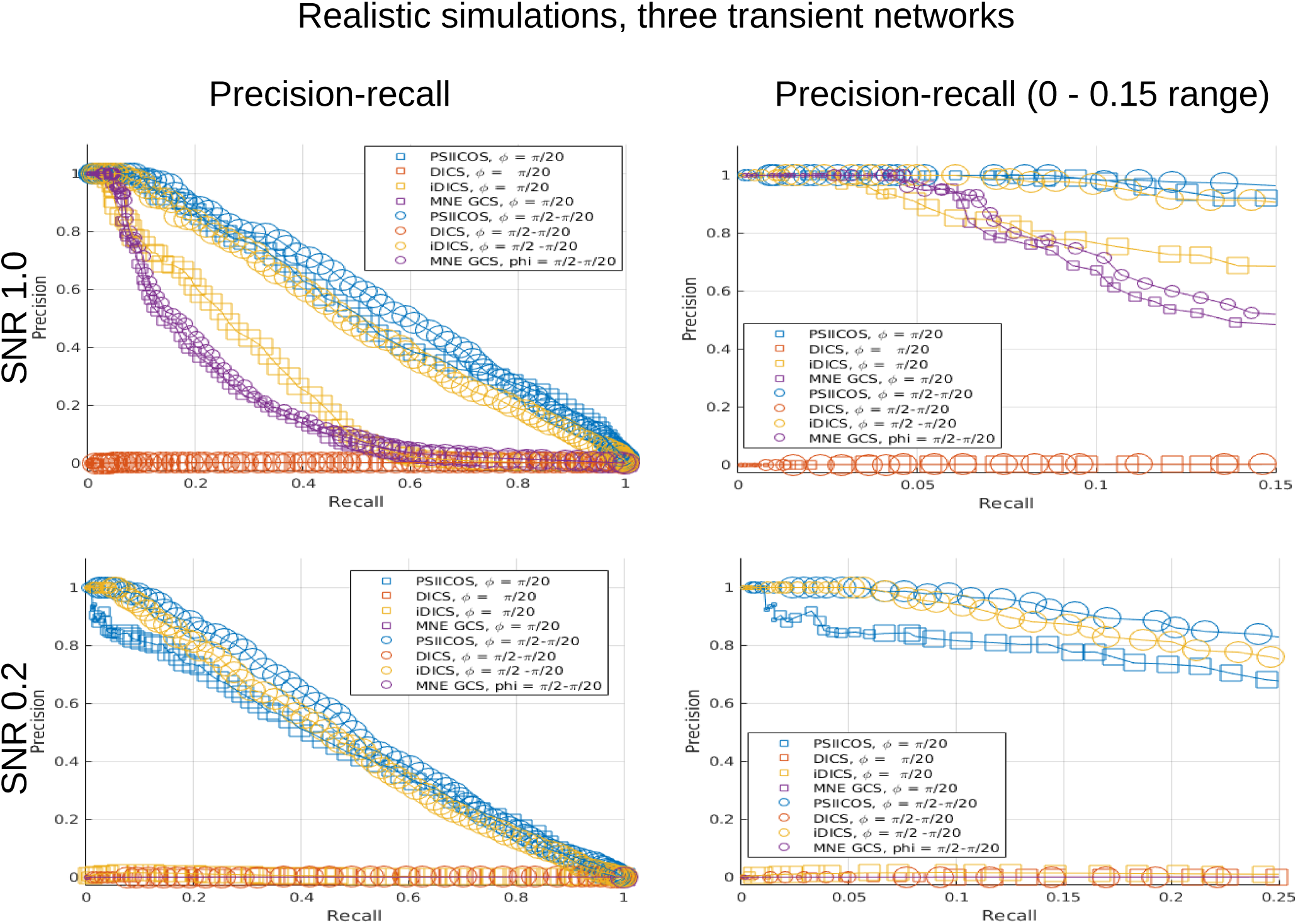
Comparative analysis of PSIICOS against the other three methods in the task of detecting the simulated three simultaneously active networks. The left half of the plots shows the entire precision-recall plot and the right panel focuses on the 0-0.15 range.

As demonstrated by our simulations the proposed approach allows to achieve superior performance in detecting of networks irrespective of the phase lag value. To illustrate this we applied the proposed technique to the analysis the simulated 3 networks data for a grid of with phase lag values. We then quantify the performance as the area under the precision-recall curve and illustrate the obtained results for using the imaginary part only, the projected real-part only and the total projected cross-spectral matrix. As we can see from figure [11] for the close to *π/*2 phase lag values the information about the interacting sources is primarily contained in the imaginary part of the cross-spectrum. Thus, the use of the imaginary part only allows to achieve a high performance which deteriorates as the phase lag gets reduced. On the contrary, for close to zero phase lags the coupling information is primarily present in the real part. The real part is contaminated by the SL component that can be efficiently removed by the proposed projection procedure. Then, the real part can be used for detection of networks with close to zero phase lag. Simultaneous use of both real and imaginary components allows to achieve a uniform performance over various phase lag values.

**Figure 11:**
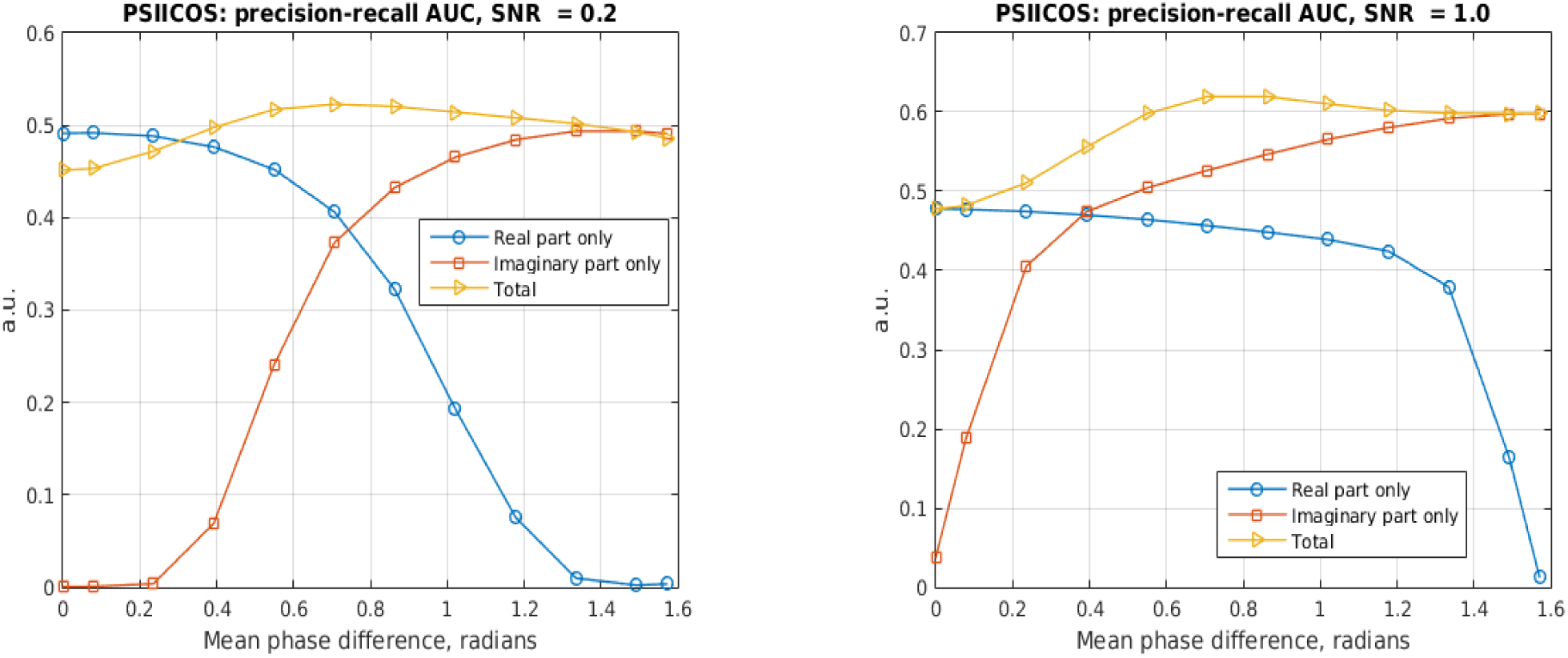
For the close to *π/*2 phase lag values the information about the interacting sources is primarily contained in the imaginary part of the cross-spectrum. Thus, the use of the imaginary part only allows to achieve a high performance which deteriorates as the phase lag gets reduced. On the contrary, for close to zero phase lags the coupling information is primarily present in the real part. The real part is contaminated by the SL component that can be efficiently removed by the proposed projection procedure. Then, the real part can be used for detection of networks with close to zero phase lag. Simultaneous use of both real and imaginary components allows to achieve a uniform performance over various phase lag values.

## 4 Real data results

To assess performance of the algorithm on real data we took recorded MEG data for 3 subjects performing volitional finger movements each 2-3 seconds. Movement onset was registered via accelerometer attached to the left hand index finger for each subject. The maximum of accelerometer response was taken as a movement event. We compared seed-based connectivity during movement preparation (-0.6 to -0.2 seconds relative to movement) to poststim connectivity averaged on 0 to 0.4 seconds time interval. Seed was placed in the left M1 area and plotted as a red sphere on fig [12].

**Figure 12:**
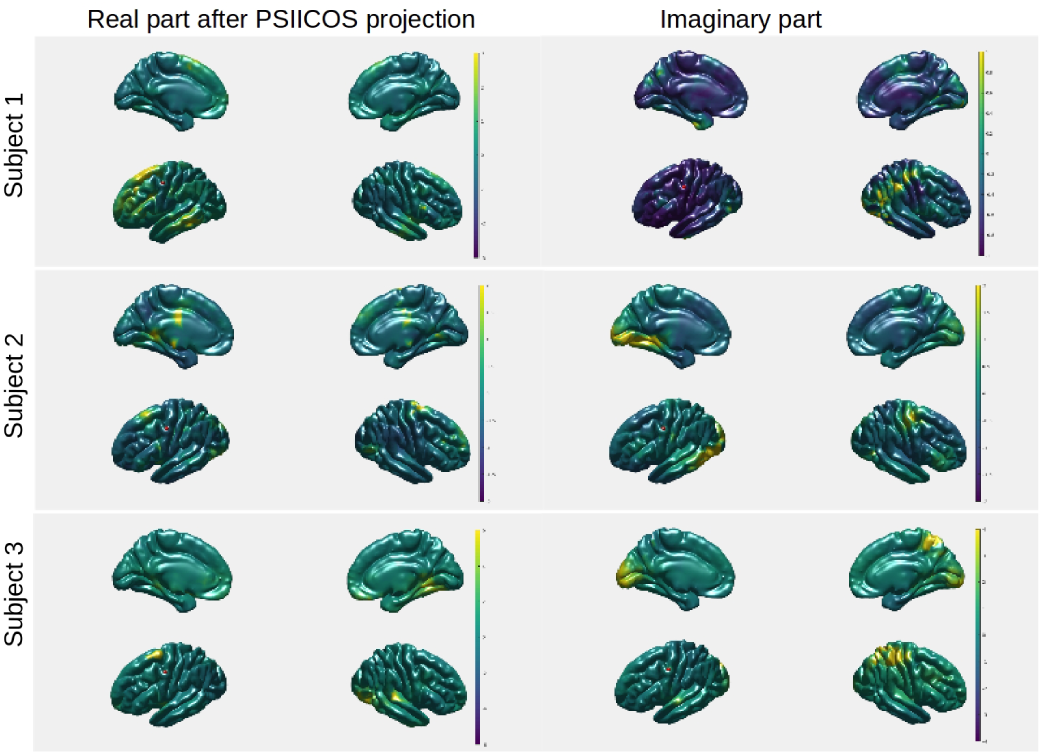
Prestim -postsim relative difference in seed-based connectivity for real projected cross-spectrum vs imaginary cross-spectrum on 3 subjects in volitional left finger movements paradigm. Seed is located in the left M1 hand-representation area (red sphere on plots). For each of three subjects we′ve found coherence between seed location and premotor area using projected real part of the cross-spectrum (left column). This effect wasn′t observed for an imaginary part of cross-spectrum (right column).

## 5 Discussion

We have described a novel method for detection of within frequency coupling based on the non-invasive MEG and EEG data. The main hallmark of the method is its nearly uniform sensitivity over the entire range of phase-lag values. This is achieved by the proposed projection procedure that operates in the product space of cross-spectral matrices and allows to efficiently suppress the spatial leakage (SL)contribution to the real part of the cross-spectral matrix. It turns out that while the subspace modulated by the real part of the source space cross-spectrum and the subspace that encapsulates the SL power overlap it is quite straightforward to build a spatial projector to banish 95% of the SL power and yet remain sensitive to majority of zero-phase coupled sources. The projector is built using the SVD of the SL "topographies" to concentrate maximum of the SL energy in a subspace of a smallest rank.

We have conducted an extensive investigation of the proposed technique performance and compared it to the DICS, the imaginary DICS and the Geometric Correction Scheme (GCS) techniques in the realistic simulations setting. The PSIICOS technique consistently outperformed the other methods in both Monte-Carlo and fixed configuration three networks study under a wide range of noisy conditions and phase-lag values.

Numerous techniques have recently emerged to non-invasively detect functionally coupled networks. Should we have access to the actual signals reflecting the activity of each individual network′s node we could use the the coherence function that reflects linear (from the LTI theory point of view) relationship between the signals. However, the activity of cortical generators as measured by the non-invasive sensors is only available in the form of a mixture of activation signals of multiple sources and thus the direct use of sensor signals leads to erroneous detection when the spatial leakage effect masks the true functional coupling. To overcome this problem Guido Nolte suggested the use of the imaginary part of the cross-spectrum as a sensor-space statistics insensitive to the spatial leakage effects. This triggered the appearance of the whole range of methods exploiting the imaginary part of the sensor-space cross-spectrum. While some of them do render an advantage over the basic imCoh technique they all are unable to detect zero phase coupled networks as the imaginary part of the sensor-space coherence is not modulated by the real part of the source-space cross-spectrum that carries the information about the actual zero-phase coupling. For low phase lags the SNR of the imaginary part of the sensor-space cross-spectrum is not sufficient to produce a reliable detection, see figure [11]. However, the use of the raw real part is not possible due to the spatial leakage effect. As we have shown in our simulations the use of an inverse operator to transform the sensor-space cross-spectrum as suggested by the DICS technique, and subsequent use of both real and imaginary parts of the resultant source-space coherence fails to deliver a reasonable performance for the realistic SNR scenario.

The geometric correction scheme (GCS) Weins et al suggests the use of the seed node topography combined with the inverse operator row corresponding to the probed source location to remove the spatial leakage effect. The use of this approach was demonstrated on analysis of envelope functional coupling and compared against the timeseries orthogonalization techniques that relate closely to the imaginary coherence based techniques when applied to analysis of the linear or phase functional coupling.

The approach proposed in this paper is most closely related to the GCS by Weins but instead of using the seed source topography and removing the SL effect of only the single (seed) source the PSSICOS approach operates in the product space and creates the projector that takes into account the contributions of all sources to the SL effect. The use of the SVD allows to build a projector that efficiently utilizes the degrees of freedom and concentrates the largest amount of the undesired SL power in the subspace of the smallest dimension. The projection operation of PSIICOS is applied to the sensor signals cross-spectral matrix and allows to free it from the SL effect. Thus, unlike the GCS the PSIICOS allows to visualize the dynamics of interaction dependent cross-spectrum in the sensor space and provides an opportunity for subsequent sensor-space analysis (not pursued in this paper). For example, given the growing utility of the machine learning approaches in analysis of the neuroimaging data the projection operation forming the foundation of PSIICOS allows to obtain the features reflecting the true connectivity yet in a relatively compact product space of sensor signals. At the same time, the GCS technique applied at the stage of extracting the source signals (corrected for the seed SL effect) allows to compute arbitrary non-linear FC statistics (envelope correlation, phase-amplitude coupling, etc) which is not possible with the proposed technique.

The proposed linear algebra based point of view allows to interpret the task of estimation of the sensor-space cross-spectrum generative model parameters as a standard underdetermined linear regression problem. This approach shows a clear way to introduce much needed priors into the connectivity estimation task. The priors can be extracted from the DTI results and represented using the probabilistic distribution to be then naturally exploited within the Bayesian paradigm. Simpler priors based on the sparsity can also be utilized and an approach similar to that of A. Gramfort exploiting the mixed fractional norms of the obtained solution can be utilized to obtain sparse solutions explaining the observed sensor space cross-spectrum.

Following the parametric estimation path it is possible to apply the extension of dipole fitting techniques including the modified RAP-MUSIC algorithm to the SL free cross-spectral matrix. In fact, describe the used of the MUSIC approach for analysis of the imaginary part of the cross-spectrum using the MUSIC metrics. Unfortunately, because of the use of the imaginary part of the cross-spectrum the proposed approach fails to be sensitive to zero phase-lag coupled networks. Also, the computational time of the described procedure is high and the authors resort to the two stage procedure to avoid the scan over *N* ^2^ source pairs.

In the current vectorized implementation in the Matlab environment, computing the projection matrix for the cortical source model with 3000 nodes takes under one second time and has to be done once per subject, given the sensor positions are fixed or the data are corrected for the movements. Then, the scan over 3000*×* 3000 sites takes less than a second. Therefore, the proposed approach is computationally efficient and allows to perform bootstrap analysis for investigation of the stability of the observed networks as it was done in Darvas et al. [2005] for a more standard dipole fitting task.

## 6 Acknowledgement

The authors would like to thank the reviewers for the valuable comments that made this manuscript a better read for a broader audience. The study has been funded by the Russian Academic Excellence Project ′5-100′ and by the RFBR grant 14-02-00917.

